# Consistency of social interactions in sooty mangabeys and chimpanzees

**DOI:** 10.1101/2020.07.10.196949

**Authors:** Alexander Mielke, Anna Preis, Liran Samuni, Jan F. Gogarten, Jack Lester, Catherine Crockford, Roman M. Wittig

**Affiliations:** Primate Models for Behavioural Evolution Lab, University of Oxford, UK; Taï Chimpanzee Project, Centre Suisse de Recherches Scientifiques en Côte d’Ivoire, Abidjan, Côte d’Ivoire; Wild Chimpanzee Foundation; Department of Human Evolutionary Biology, Harvard University, USA; Robert Koch Institute, P3: “Epidemiology of Highly Pathogenic Microorganisms”, Berlin, Germany; Max-Planck-Institute for Evolutionary Anthropology, Leipzig, Germany

## Abstract

Predictability of social interactions can be an important measure for the social complexity of an animal group. Predictability is partially dependent on how consistent interaction patterns are over time: does the behaviour on one day explain the behaviour on another? We developed a consistency measure that serves two functions: detecting which interaction types in a data set are so inconsistent that including them in further analyses risks introducing unexplained error; and comparatively quantifying differences in consistency within and between animal groups. We applied the consistency measure to simulated data and field data for one group of sooty mangabeys (*Cercocebus atys atys*) and to groups of Western chimpanzees (*Pan troglodytes verus*) in the Taï National Park, Côte d’Ivoire, to test its properties and compare consistency across groups. The consistency measures successfully identified interaction types whose low internal consistency would likely create analytical problems. Species-level differences in consistency were less pronounced than differences within groups: in all groups, aggression and dominance interactions were the most consistent, followed by grooming; spatial proximity at different levels was much less consistent than directed interactions. Our consistency measure can facilitate decision making of researchers wondering whether to include interaction types in their analyses or social networks and allows us to compare interaction types within and between species regarding their predictability.

## INTRODUCTION

Animals living in permanent social groups must decide when and how to interact with group members, and their ability to make appropriate choices has potential fitness implications (Shettleworth, 2009). The evolution of species’ cognitive apparatus is a response to selection pressures imposed by the complexity of their environment, including the social system they live in (Byrne & Whiten, 1989; Humphrey, 1976; Jolly, 1966). This hypothesis assumes that animals in more “complex” social systems must integrate more social information to out-compete others (Byrne & Whiten, 1989). However, it is unclear how to quantify social information, even though various indices have been proposed (Bergman & Beehner, 2015; Fischer et al., 2017).

One way to operationalize social complexity is as the amount of information necessary to successfully predict future states within a system (Flack, 2012; Sambrook & Whiten, 1997). Measures of interaction predictability on an individual level in group-living species would facilitate examinations of factors driving evolution of complex decision-making (Aureli & Schino, 2019; Dunbar & Shultz, 2010).

Consistency of partner choice across time, i.e. repeatedly choosing to interact with the same individual in the same way, enhances the predictability of future outcomes (Kalbitz et al., 2016; Koski et al., 2012; Moscovice et al., 2017; Silk et al., 2006). For example, in steep linear dominance hierarchies, a single interaction per dyad contains enough information to predict future dyadic contests (Guillermo Paz-Y-Miño et al., 2004; Oliveira et al., 1998; Sánchez-Tójar et al., 2018). Low consistency can be the result of an unpredictable distribution of social interactions, frequent changes in relationships over time, or the presence of various mediating factors, all challenges that might necessitate an increased need for cognitive flexibility (Barrett et al., 2002).

Assessing predictability is complicated by the fact that we work with incomplete data, as recording every interaction taking place in an animal group is not practicable. Many behavioural studies depend on aggregated distributions of interactions over time: we take, for example, a one-year period and calculate how many interactions were observed on an individual and dyadic level. These distributions are used either as dependent or independent variables, to create networks, or to create relationship indices. The fundamental assumption is that the data accurately reflect what individuals were doing during the study period and that patterns are consistent; the “real” distribution of interactions is unknown (Farine & Strandburg-Peshkin, 2015; Kasper & Voelkl, 2009; Whitehead, 2008). However, if data are sparse, estimate errors are increased and robustness of the resulting distribution reduced (Lusseau et al., 2008; Shizuka & Farine, 2016). Working with measures that are not accurate representations of the underlying distribution can create misleading results (Davis et al., 2018). This problem is exacerbated when already sparse datasets are cut into shorter time intervals (e.g. 6-month blocks), a common practice in animal behaviour studies. What constitutes ‘enough’ data can vary depending on the consistency of partner choice (Sánchez-Tójar et al., 2018; Whitehead, 2008). For many researchers, it is difficult to assess whether they have collected enough data to include an interaction type into their analyses. Here, we propose a shorthand.

In the present study, we develop a consistency measure that serves two functions: 1) allowing researchers to gauge whether they have collected enough data to warrant the inclusion of an interaction type in their analyses, in a social network, or when creating relationship indices. 2) Compare predictability of interaction types within, between, and among species. Consistency should be high if individuals choose the same partners for the same interaction type independent of when they are observed, and observing an individual at one point in time allows for accurate predictions of its future behaviour. Low consistency can arise if individuals show weak partner preference or preference changes over time, or if insufficient data are available.

To explore how consistency can be used to compare social groups with different structure and organisation, we first use simulations of datasets with different properties. We subsequently apply the consistency measure to data from two Western chimpanzee (*Pan troglodytes verus*) communities and one sooty mangabey (*Cercocebus atys atys*) community living sympatrically in the Taï National Park, Côte d’Ivoire (Mielke et al., 2017, 2018). These species represent two well-studied, quite different primate social systems. Sooty mangabeys have philopatric females who form linear, despotic, stable matrilineal hierarchies (Mielke et al., 2017, 2018; Range, 2006; Range & Noë, 2002). All mangabey directed social interactions are expected to show high consistency, as they should be strongly influenced by stable parameters, especially kinship, dominance rank, and sex (Range & Noë, 2002). Association patterns in this species are nearly random (Mielke et al., 2020), so we predict low consistency for spatial interaction types. Chimpanzees at Taï are similar in that they have been shown to have stable grooming, aggression, and association patterns in both sexes. However, in contrast to the mangabeys, aggression is not exclusively determined by dominance hierarchy (Wittig & Boesch, 2003), and we have previously described rank changes in both sexes in the study period (Mielke et al., 2019; Preis et al., 2019). Rank uncertainty and the variation in partner availability due to fission fusion dynamics in chimpanzees lead us to predict that chimpanzee interactions are less consistent than mangabey interactions. We developed the consistency measure with two aims: a) to identify interaction types where data distributions are likely unreliable due to insufficient data; and b) to draw comparisons between chimpanzees and mangabeys, and identify different interaction types within species as it relates to their consistency.

## METHODS

### Consistency measure

To quantify consistency in an interaction type, we organised the data by collection days. Each observation day is randomly assigned to one of two datasets of equal size (Sánchez-Tójar et al., 2018). For each of the two resulting datasets, we calculated the dyadic interaction rates per observation hour in each of the halves and calculate the non-parametric Spearman correlation between the two distributions (see Fig. 1 for procedure). This allowed us to estimate how well variation in one half of the dataset predicts variation in the other half. We performed 100 iterations, with the median correlation coefficient constituting our measure of consistency for the full dataset.

**Fig. 1:**
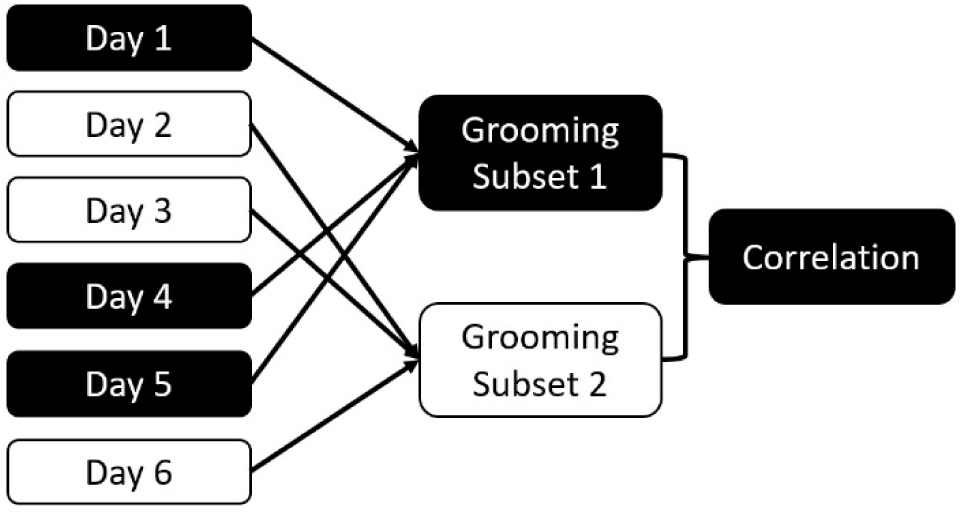
Schema of the consistency measure. Data are randomly divided into two subsets based on the collection day. Dyadic interactions for all dyads are aggregated for each subset. The two subsets are correlated. This process is repeated 100 times to calculate the consistency of the overall dataset for this interaction type.

The overall correlation between halves of the dataset is likely dependent on the data collection effort and community size, making it difficult to compare communities and interaction types. To mitigate this challenge, we developed a standardised version of the consistency measure (Fig. 2) by repeatedly selecting subsets of the data that differ in length and the amount of data included, followed by randomly selecting a start date and duration for the period following that date. We tested the consistency for this period for each interaction type, marking how many interactions per dyad the subset contained. For example, 10 individuals form 45 dyads; if we collect 180 aggressive events, we have a mean of 4 interactions/dyad. We then collate the consistency of all datasets with the same number of interactions per dyad – *e.g*., we could have 100 consistency values based on datasets that contain 3 interactions per dyad, 120 based on 4 interactions per dyad, and so on. For each interaction per dyad value, we plot and report the median of the consistency values.

**Fig. 2:**
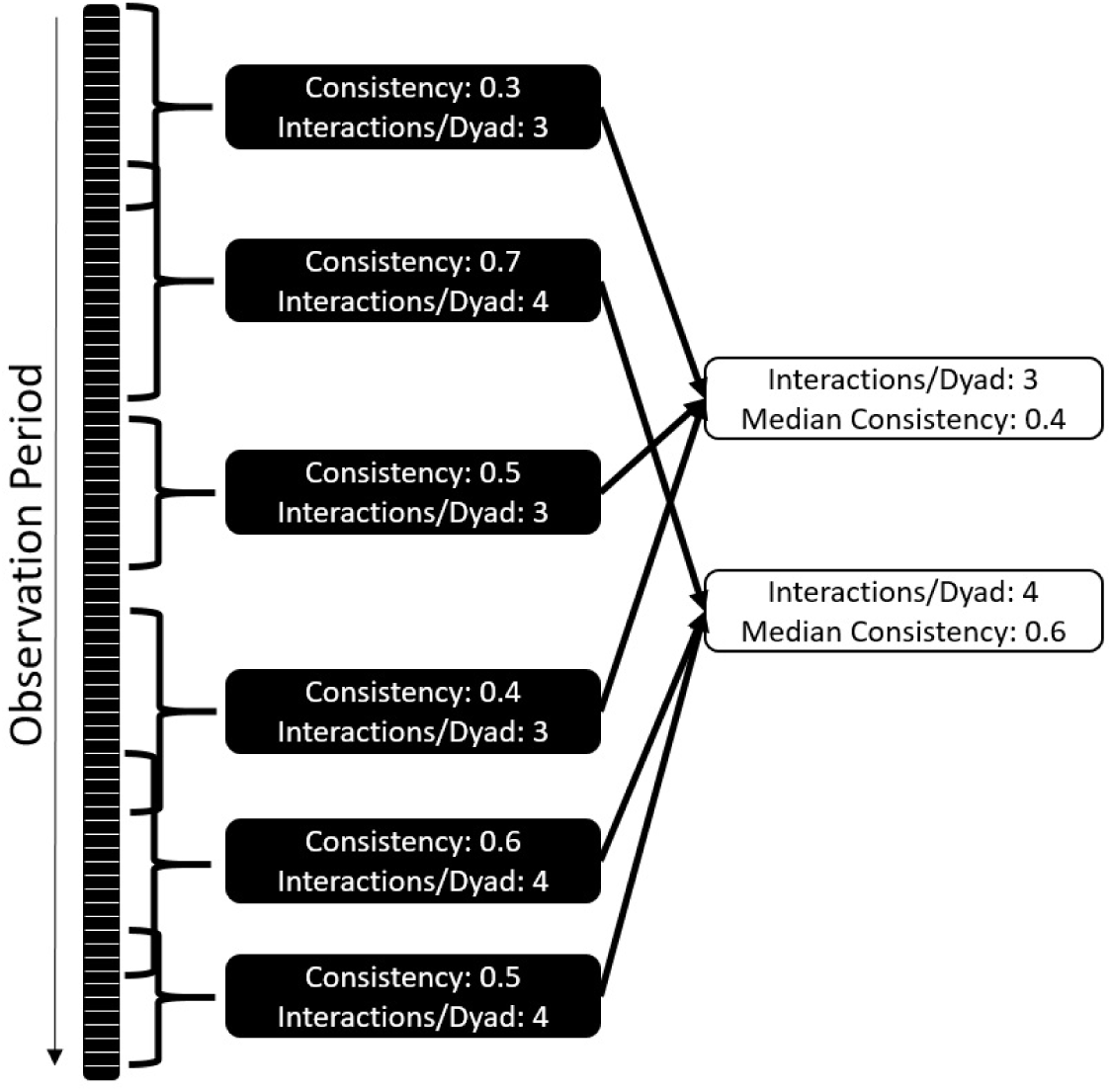
Standardisation of consistency as a comparative measure: each dataset is randomly cut into shorter time windows of changing size and starting point. Consistency and amount of interactions per dyad are established. The median of consistencies across different interactions per dyad values are established. The comparative value is the number of interactions per dyad where the consistencies cross 0.5.

This approach allows a systematic comparison of both frequent and infrequent interaction types, *i.e*., datasets of different sizes. Analyses comprising differing group sizes are possible because we compare the behaviour of datasets that contain the same number of interactions per dyad. As a standardised consistency measure, we report the number of interactions per dyad needed to get a median consistency value of 0.5; although this value has no strong biological justification, in simulations it was reliable in distinguishing interactions types that were consistent from those that had insufficient data or were inconsistent. This measure is largely independent of data density and community size, and produces an interpretable result: how many interactions between two group members does an individual need to observe to reliably predict future interactions? Fewer interactions per dyad and a smaller standard deviation of values indicate higher consistency in partner choice and thus higher predictability. Larger numbers of interactions per dyad and a large standard deviation indicate that interaction patterns are harder to predict. This can be the case if either the partner choice is less deterministic for the interaction type, or the choice patterns change throughout the study period.

### Simulations

All described analyses were conducted in R 4.0.0 (R Development Core Team & R Core Team, 2020). Scripts can be found in the associated GitHub repository. We explored the impact of different group sizes, data densities, repeatability of partner choice, and changes in underlying relationships on our consistency measure using simulated datasets. We then explore whether it can be used to compare consistency across communities of different sizes. We tested whether our consistency metric is high when individuals regularly choose the same partners for the same interaction type. We also tested whether low consistency arises when individuals show weak partner preference or when preference changes over time. To test how our consistency measure performed under different conditions, and how to interpret different results, we simulated datasets with different group sizes; numbers of interactions per individual; data collection density; stereotypy of partner choice; and consistency of partner choice over time, mirroring interaction data as it could be collected in different social animal species.

Specifically, we created datasets for 10, 15, and 20 individuals in a community, for one nonspecific interaction type over a simulated period of one year. We randomly assigned each individual between 1 and 10 interactions per day, and for each interaction the partner was chosen from a random chosen subset of group members (to simulate animal groups, in which not all group members are always physically available). To simulate different underlying probability distributions of who interacts with whom, each dyad was assigned a random likelihood to interact with each other, with three different stereotypy levels: “high certainty” (each individual has strong preference for a few group members, always chosen those when they are available), “medium certainty” (each individual prefers several group members, but can also choose non-preferred partners), and “low certainty” (the likelihood of choosing any partner is relatively equal). Based on these dyadic values, one of the individuals in the “party” was selected as interaction partner. We explored three conditions concerning the consistency of individuals’ choice: in the first condition, dyadic preferences remained the same throughout. In the second condition, to simulate changes in interaction patterns, all likelihoods of partner choice were reversed halfway through data collection, so dyads with a 0.95 likelihood of interacting in the first half had a 0.05 likelihood of interacting in the second half of data collection. In the third condition, partner choice was completely random, which should lead to an even distribution of interactions between all group members over the whole time.

Following this procedure, we created 108 simulated datasets (three each for every combination of number of individuals, level of stereotypy, and consistency condition) that contained all interactions for all group members for each day of the data collection period. Subsequently, we simulated differences in data collection effort (Davis, Crofoot, & Farine, 2018): for each day of the sampling period, one individual was chosen as the “focal” individual whose data were retained, as would be the case in most animal datasets. We assumed a twelve-hour observation period per focal day, to calculate interaction rates. Then we simulated that data collection took place every day, 66% of days, or every third day (33%), to test the impact of low data collection density on the consistency measure. We therefore retained 324 simulated datasets with different properties. For each of these, the proposed consistency measure – randomly selecting half of the dataset and correlating interaction rates of dyads with those of the other half, as well as repeating this procedure with subsets of the data – was carried out 100 times.

### Data Collection

Behavioural data were collected in Taï National Park, Côte d’Ivoire (Wittig & Boesch, 2019) from October 2013 to July 2015 for the chimpanzees and January 2014 to September 2015 for the mangabeys, using half- and full-day continuous focal animal sampling (Altmann, 1974) for the chimpanzees, and half-day and one-hour focal animal sampling for the mangabeys. Scripts and data can be found in the associated GitHub repository. Trained observers and field assistants recorded all social interactions of adult male and female chimpanzees (above 12 years of age) in the “South” and “East” communities and adult (above 4.5 years) sooty mangabeys. This resulted in 6441h of focal observations in South community, 5668h for East community, and 2259h for the mangabey community. We included adult individuals of both sexes in all three communities for whom sufficient focal data (at least 50 social interactions observed as focal individual) were available and who were present for at least 80% of the study period (South: 5 males, 7 females; East: 5 males, 7 females; mangabeys: 6 males, 17 females).

From the behavioural data, for each dyad, we extracted the duration of grooming sent, resting or foraging in less than 1m distance from the partner (“body contact”: used as a continuous measure with duration in the chimpanzees and an event variable in the mangabeys), resting or foraging as nearest neighbour between 1m and 3m distance (“proximity”), and both contact and noncontact aggressive interactions with one clear recipient (Preis et al., 2018). For the chimpanzee communities, we included food sharing (Samuni et al., 2018), which was not regularly observed in the mangabeys. We used pant grunt vocalisations in chimpanzees and feeding supplants in mangabeys as additional interaction types. Mutual interactions were coded as interactions given in both directions. We treated body contact and proximity as ‘interaction types’ with the assumption that both individuals have to show sufficient tolerance to allow the other one to remain close. Body contact and proximity were only counted if no other interaction took place within 5min before or after to ensure independence of data points. We included grooming, contact aggression, noncontact aggression, pant grunts/supplants, and food sharing as directional variables, with the distribution of interactions given from each individual to every other. For the two spatial proximity measures, data were considered non-directional and symmetrical. Interaction distributions were standardised by focal observation time, with observation time calculated by adding the total observation times of A and B. Spatial proximity and food sharing in the chimpanzees were collected by a subset of observers and were standardised based on the focal observation time provided by those observers. Scripts to create the consistency measure and plots can be found in the Supplementary.

## RESULTS

### Simulations

We sought a consistency measure that can identify differences in stereotypy of partner choice and changes in interaction preference, while being independent of group size and data collection effort. Our standardised consistency measure performed the same, independent of community size (Fig. 3.1). Meanwhile, datasets of different data collection density followed the same trajectory, but lower data density was characterized by lower overall consistency. The overall consistency cannot be interpreted alone, as it is highly dependent on group size and data collection effort. This is consistent with simulations showing that social network data becomes unreliable if data density per dyad sinks below a certain level (Whitehead, 2008).

**Figure 3:**
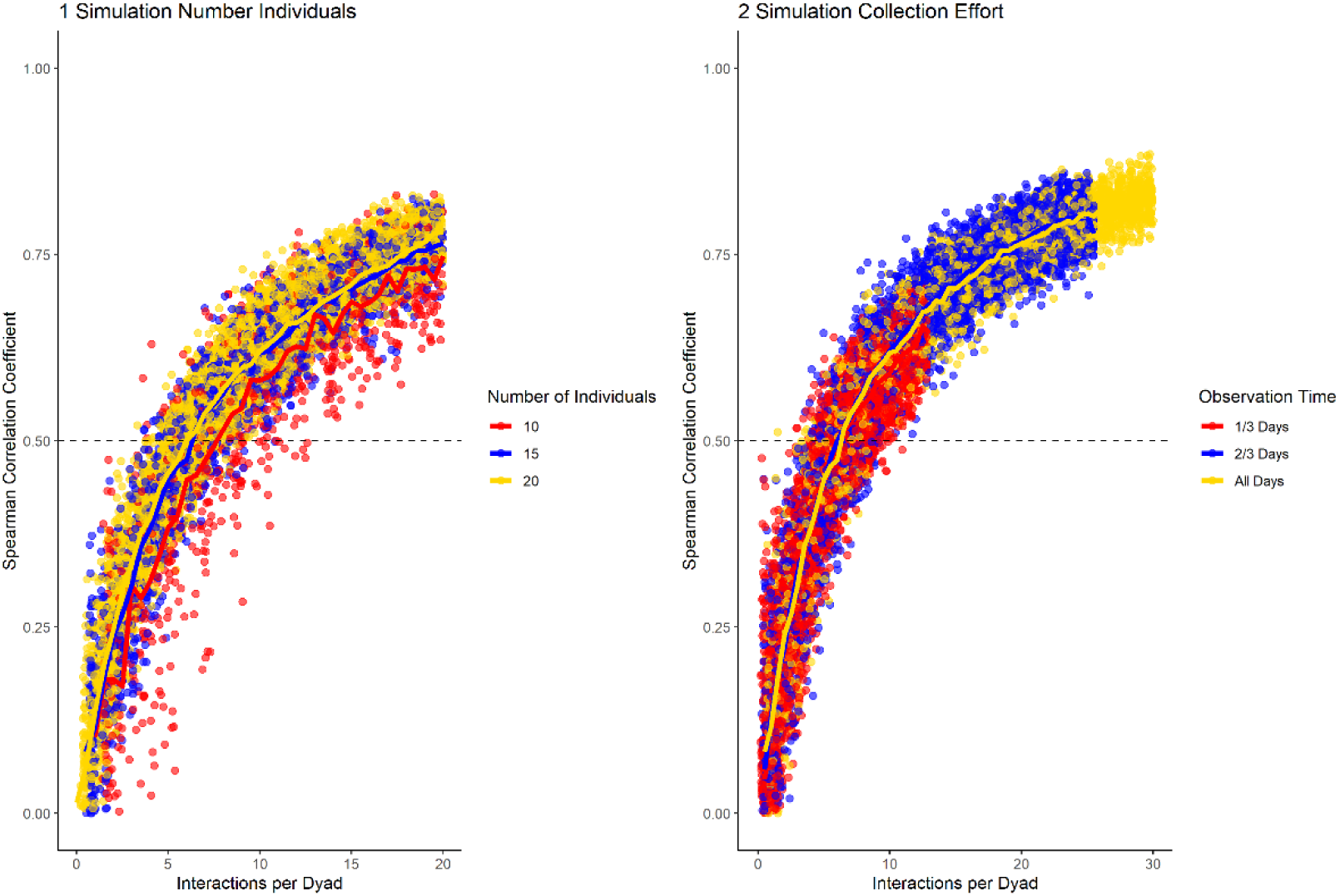
Results of the data simulation with varying group size (1) and data collection density (2), while having medium stereotypy of partner preference and no preference changes throughout the dataset. Horizontal line marks a correlation of halves of 0.5. The number of interactions per dyad allows to compare datasets of different density and number of individuals.

To test how the stereotypy of partner choice influenced the consistency measure, we present the results for the three different conditions (high, medium, low certainty) in datasets containing 20 group members, with 100% data density, and no changes in preference throughout the sampling period (Fig. 4). Our results showed that the consistency measure differentiated between the conditions, using the slope at which the chosen cut-off value is reached. If partner choice is highly stereotypical, a small number of interactions was sufficient to predict partner choice in half of the data with that of the other half; with increasing uncertainty, more interactions per dyad were necessary. For low certainty of partner choice, more than the number of simulated interactions would have been necessary to reach the cut-off of 0.5.

**Figure 2:**
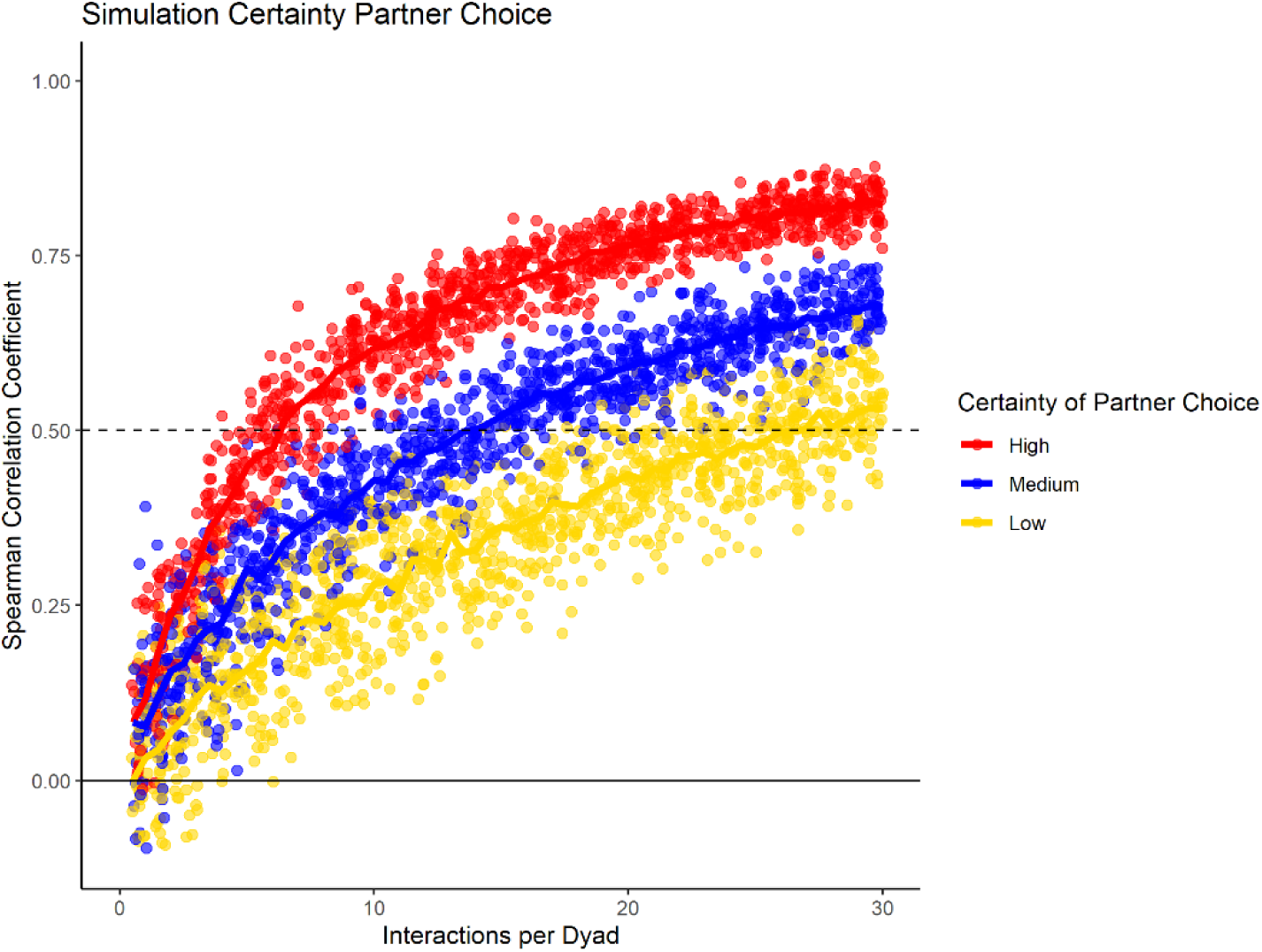
Data simulation of varying stereotypy of partner choice, while having consistent group size, data density, and no preference changes throughout the dataset.

Last, we investigated how changes in partner preference over the study period would influence the consistency measure in a dataset with 20 individuals, with 100% data density, and high stereotypy of partner choice. Here, we compare three conditions: one where no changes took place, one where the partner preference was reversed halfway through the study, and one where partner choice was randomized. Again, we found differences in the slope whereby the consistency increased with increasing data density (Fig. 5). Additionally, the conditions could be differentiated by the spread of consistency values: when partner choice was consistent, selecting subsets of the same size at different points of the sampling period resulted in very similar consistency values. If partner choice changed throughout the sampling period, variation was much higher. Also, the consistency of the full dataset was smaller than that of some shorter subsets, with the highest levels for subsets that were roughly half the total size – mirroring our built-in change of interaction likelihood after half the ‘collection period’. As seen before, random partner choice could be identified because the consistency of the full dataset never increased above a certain threshold.

**Fig. 5:**
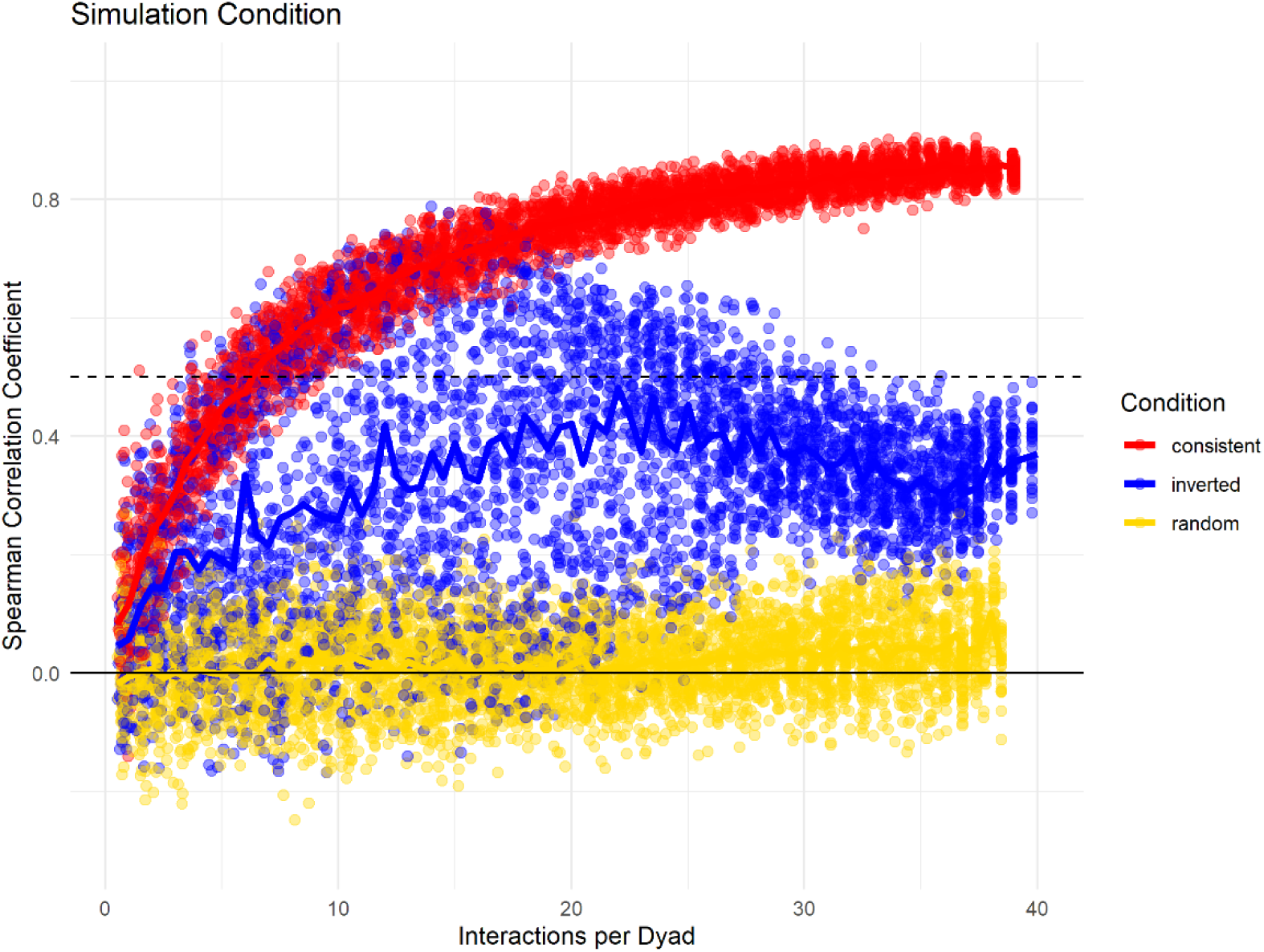
Data simulation changes potential changes of interaction distributions throughout the dataset while having consistent group size, data density, and stereotypy of dyadic preference. “Consistent Choice” indicates no changes in preferences throughout, “Inverted Choice” indicates one reversal for all dyadic preferences, while random choice indicates that all partners were chosen with the same likelihood.

In sum, based on these simulations, the consistency measure can be used to compare the predictability of interactions. Using the entire dataset, the overall consistency was heavily influenced by the amount of interactions available per dyad, and thus does not make a good comparative measure. In our simulations, even if the underlying distribution of interactions was highly stereotyped and consistent, the consistency measure remained low if few data points were available per dyad, indicating that one half of the dataset was not a good predictor of the other half due to random sampling error. Thus, if the Spearman rank correlation between halves of the same dataset does not reach *r*_*s*_ = 0.5, it is likely that not enough data have been collected to make statements about the underlying distribution of an interaction type in a population (unless that distribution is random). We therefore suggest using interaction types with an overall consistency below *r*_*s*_ = 0.5 with care or remove them from analyses where possible, as their interpretation is unclear. For all other interactions, we propose the described standardised consistency measure, the average number of interactions per dyad necessary to reach a median consistency of *r*_*s*_ = 0.5 as a good measure. Valuable information also arose from the spread of values of the repeated comparisons between halves of the dataset: if dyadic preference remained stable throughout, the consistency is relatively stable for subsets of the same size. However, if dyadic preference was not stable, the correlation between halves varies even for datasets of the same size.

### Field Data

#### Mangabeys

For the mangabeys, noncontact aggression rates (3 interactions/dyad) and supplants (3 interactions/dyad) were highly consistent, as was grooming (4.5 interactions/dyad), indicating that individuals observing a subset of interactions in the community would be able to predict future interactions (Fig. 6, 7, 8; Tab. 1). Body contact (17 interactions/dyad) was much less consistent, and proximity (being within 3m of each other) did not reach the threshold of 0.5, despite having among the highest number of data points available for any interaction type in this study. Given the trend of the graph, proximity would probably have reached the threshold if more data had been available, but this still suggests a highly inconsistent distribution across the data collection period. For contact aggression, only a small number of cases were available, and the graph did not reach the consistency threshold. In our simulations, such low values occurred when insufficient data were available to successfully approximate the underlying distributions of interactions, even in cases where the underlying distribution was highly consistent; or when distribution of interaction was random or close to random.

**Figure 6:**
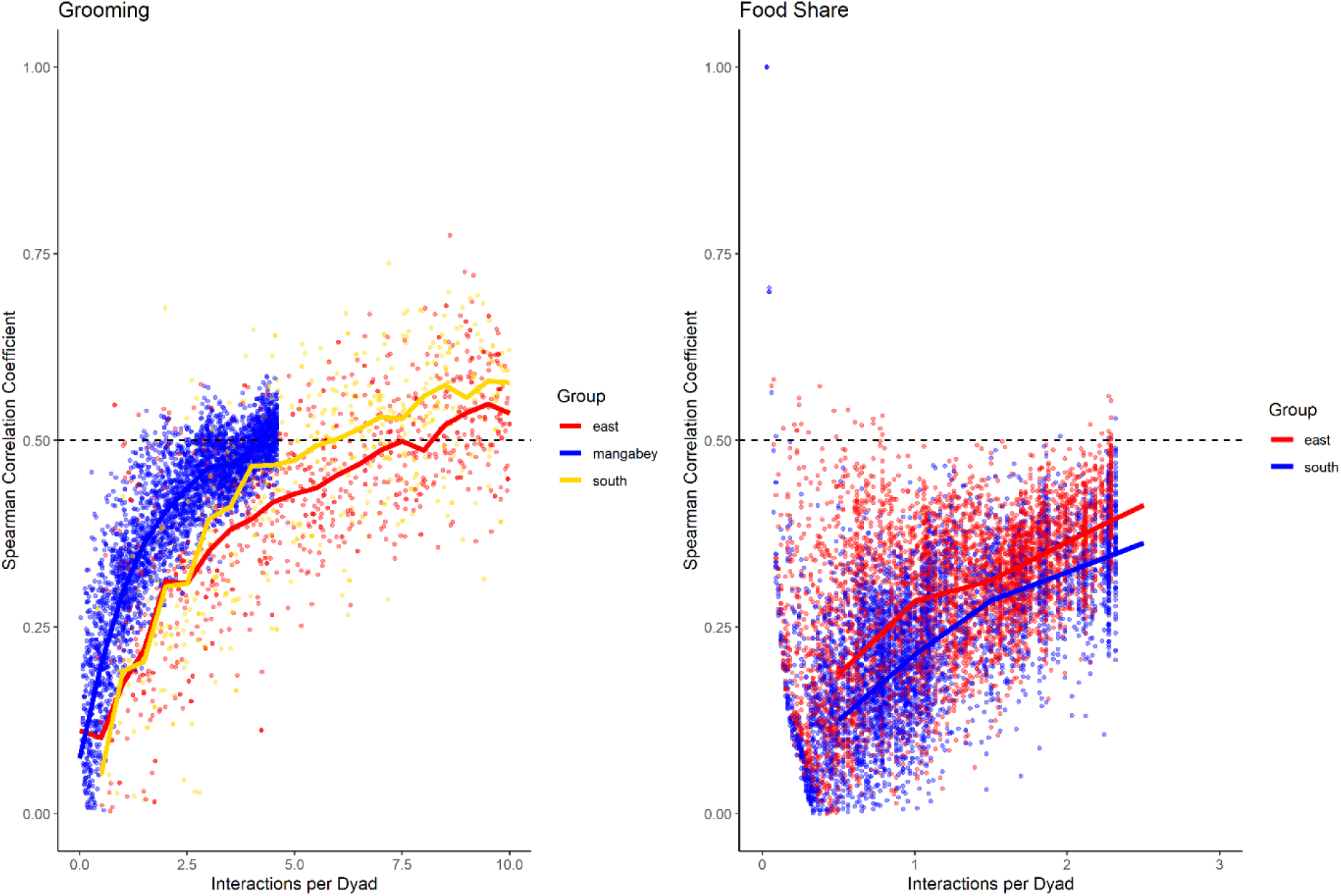
Spearman correlation between two halves of randomly selected subsets of the datasets for mangabeys (blue), East chimpanzee community (red) and South chimpanzee community (golden) for grooming and food sharing (chimpanzees only). The standardised consistency is marked by the number of interactions per dyad where the median of correlation coefficients exceeds r=0.5. If that value is reached with fewer interactions per dyad, the distribution of interaction rates is more consistent. Distributions of correlation coefficients with a large spread indicate changes in interaction preference over time.

**Figure 7:**
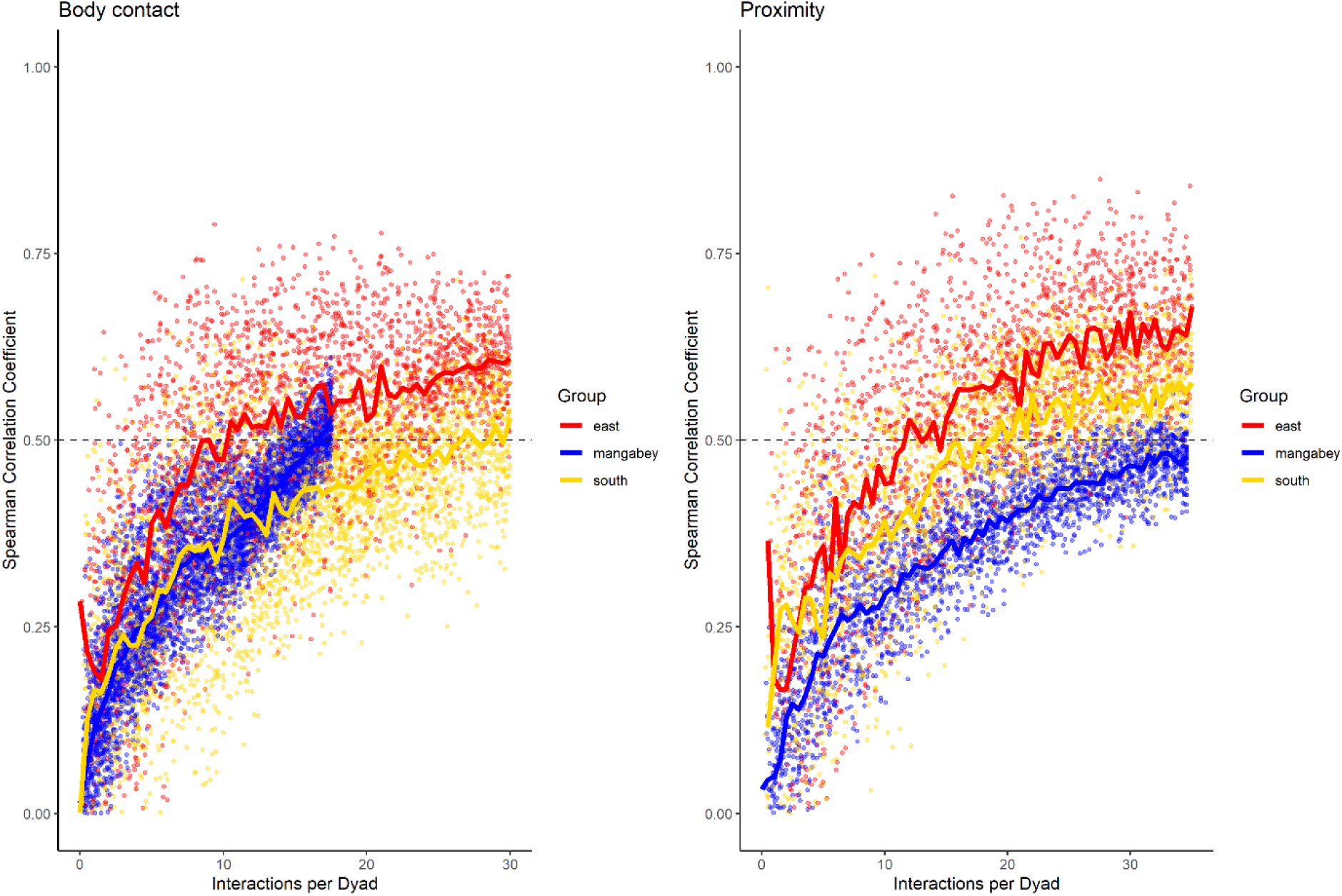
Spearman correlation between two halves of randomly selected subsets of the datasets for mangabeys (blue), East chimpanzee community (red) and South chimpanzee community (golden) for body contact and proximity. The standardised consistency is marked by the number of interactions per dyad where the median of correlation coefficients exceeds r _s_= 0.5. If that value is reached with fewer interactions per dyad, the distribution of interaction rates is more consistent. Distributions of correlation coefficients with a large spread indicate changes in interaction preference over time.

**Figure 8:**
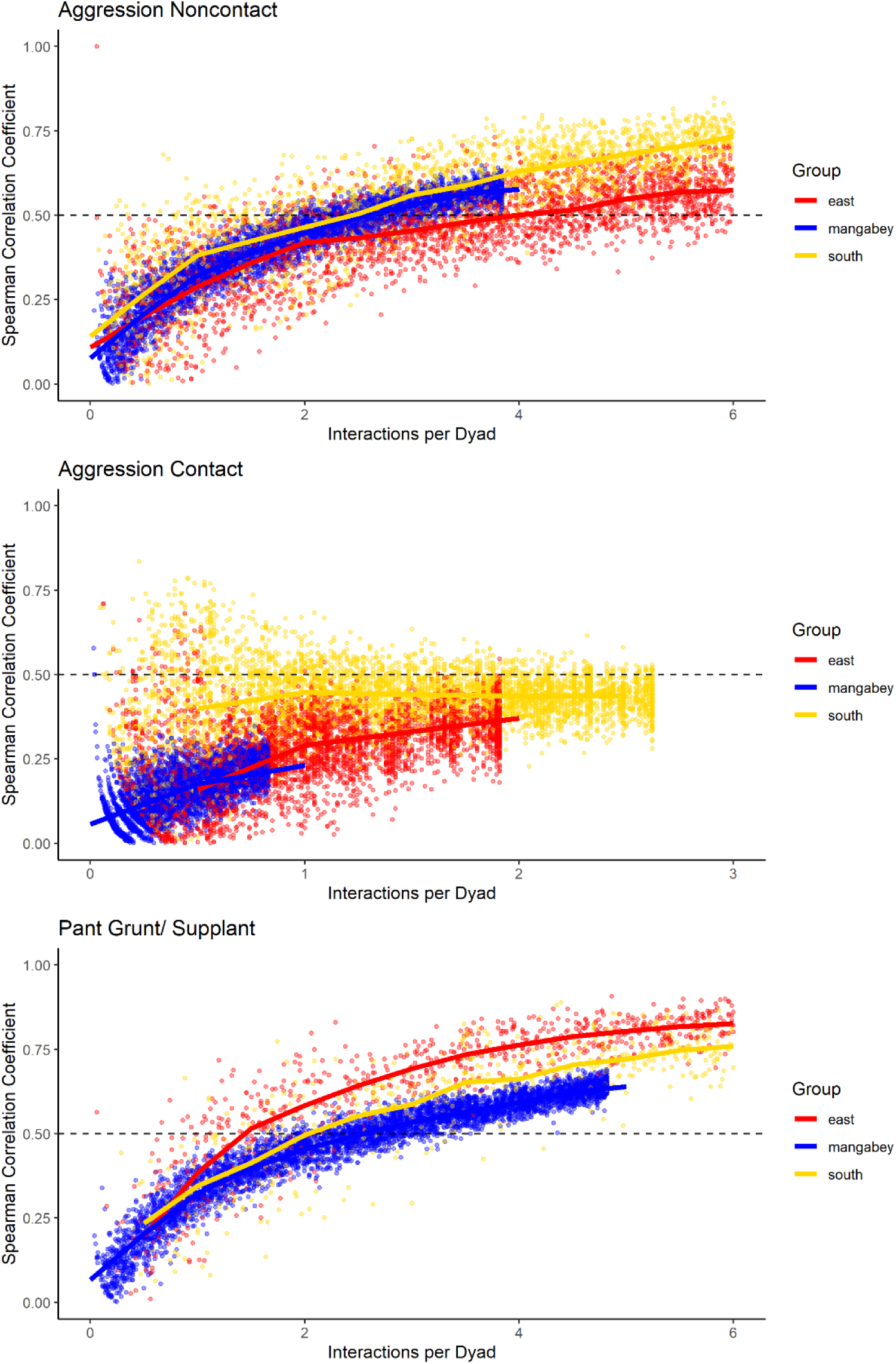
Spearman correlation between two halves of randomly selected subsets of the datasets for mangabeys (blue), East chimpanzee community (red) and South chimpanzee community (golden) for noncontact aggression, contact aggression, and pant grunts/supplants. The standardised consistency is marked by the number of interactions per dyad where the median of correlation coefficients exceeds r_s_ = 0.5. If that value is reached with fewer interactions per dyad, the distribution of interaction rates is more consistent. Distributions of correlation coefficients with a large spread indicate changes in interaction preference over time.

#### Chimpanzees

As in the mangabeys, noncontact aggression rates were highly consistent in both chimpanzee communities (Table 1), more so in South (2.5 interactions/dyad) than in East (4.0 interactions/dyad). As in the mangabeys, contact aggression occurred so infrequently that now consistent representation of the distribution existed. The larger standard deviation in the chimpanzees and wider spread of the graph compared to the mangabeys might indicate changes of aggression patterns over time. Pant grunt interactions in both communities showed the most predictable patterns (East: 1.5 interactions/dyad; South: 2.5 interactions/dyad). Grooming was less consistent than in the mangabeys (East: 8.5 interactions/dyad; South: 6.0 interactions/dyad). Body contact showed considerable variation between groups, with East (9 interactions/dyad) being the most consistent, while South (27.0 interactions/dyad) being the least consistent of the three groups. Proximity (East: 12.0 interactions/dyad; South: 19.0 interactions/dyad) was more predictable than in the mangabeys. Body contact and proximity were considerably less predictable than the directed interaction types. This indicates that in all three communities, most dyads will feed and rest in proximity with a wide variety of partners, while they direct interactions at a smaller and more stable subset of group members.

**Table 1:**
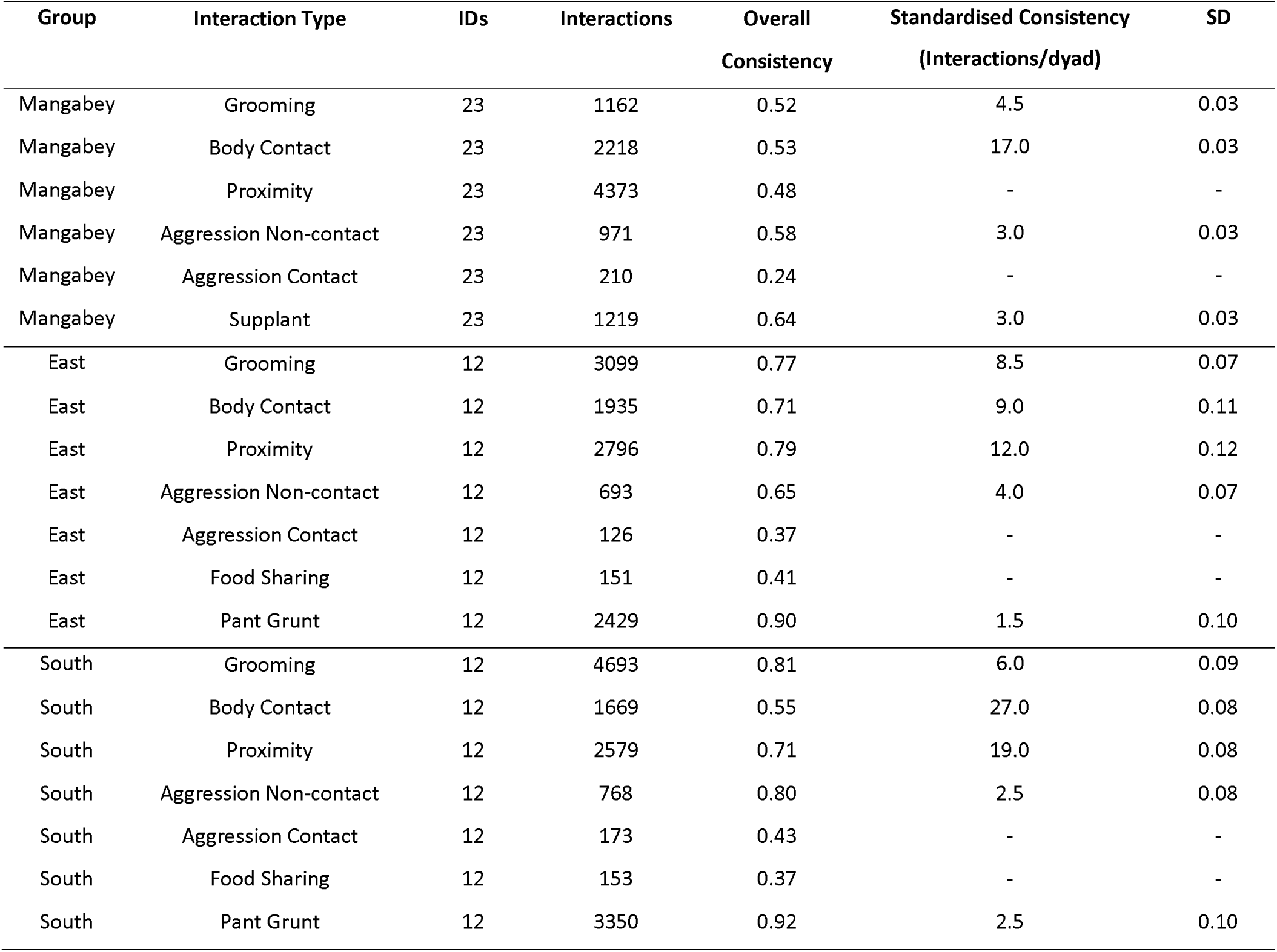
Overview of consistency scores in chimpanzee and mangabey social interactions: datasets for each interaction type and group, and the results of the consistency measures. “Overall consistency” is the median of the repeated correlation between randomly selected halves for the full dataset available for an interaction type. “Standardised Consistency” and the standard deviation are the result of the repeated random selection of halves of subsets of different lengths, with number of interactions per dyad where the median consistency exceeds r_s_= 0.5 as measure of how much information is needed to predict future interactions in a community. Interaction types were the r_s_= 0.5 was not exceeded are marked with ‘-’.

**Figure 9:**
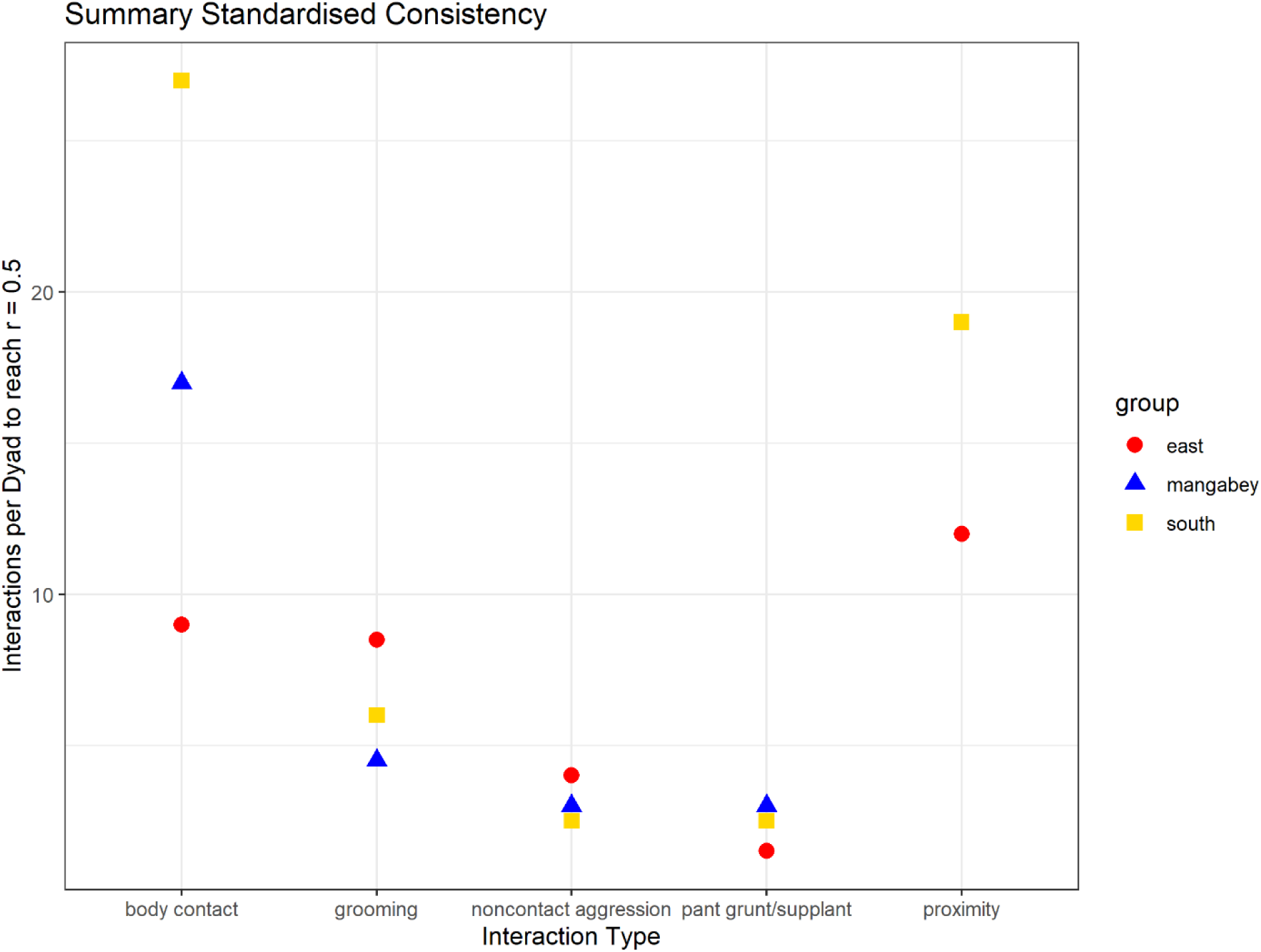
Summary of the mean number of interactions needed per dyad to reach correlations between halves of r_s_ = 0.5 (mangabeys: blue triangles, East: red points, South: golden square).

## DISCUSSION

Establishing measures of predictability of social interactions between individuals is necessary to understand the complexity of a social group from the perspective of the individual (Dunbar & Shultz, 2010; Lukas & Clutton-Brock, 2018). Here, our premise was that interactions are more predictable for participants and bystanders if interaction distributions are consistent over time. Our results showed that across communities and species, interaction types vary in predictability, indicating yet again that animal lives cannot be captured using one simplistic measure of complexity: challenges differ within and between species, and we need multi-dimensional measures to quantify where ‘complexity’ really arises.

This study introduces a consistency measure, repeatedly dividing the dataset into halves and comparing how well these predict each other, which serves two functions. Researchers can use it to find out whether they have collected sufficient data for their dataset to be internally consistent, given a community of a certain size and an interaction type with a specific diversity of partner choice (Sánchez-Tójar et al., 2018). In our sample, despite pooling 18 months of data, food sharing and contact aggression were observed at such low rates in all three communities that observing the group at a certain time point would make it impossible to predict their behaviour at another time point. We do not know whether the error bars around the observed values are biological or statistical, but they can introduce unexplained uncertainty in our subsequent analyses. We generally assume that randomly selected focal follows allow us to also make statements about interaction rates on those days on which we do not observe an individual (Altmann, 1974), but this might not be the case for rare interaction types or for interaction types that are naturally almost randomly distributed (Davis et al., 2018). If the distribution of interactions in the group is not even consistent within an interaction type in a period, correlating it with other interaction types or across periods would probably produce spurious results (Whitehead, 2008).

The standardized consistency measure allowed us to identify interaction types that needed either large or small amounts of information to predict interactions on other collection days. We used the number of interactions per dyad at which the majority of subset correlations exceeds the value *r*_*s*_ = 0.5; while the value *r*_*s*_ = 0.5 itself is arbitrary, using it across species and interaction types allows researchers to make comparative statements, and it is high enough to not fall into random variation. We did not find generalizable species differences using our consistency measure: differences within species were much more pronounced and followed the same trends between species. Chimpanzee distributions had generally larger standard deviations, potentially indicating changes in partner choice over time. Feeding supplants and pant grunts, which are used to create hierarchies in the respective species, were highly consistent, indicating generally stable hierarchies (Sánchez-Tójar et al., 2018). Consistency of aggression distributions did not vary strongly between species. Despite being the larger community, mangabey grooming interactions were generally more predictable than chimpanzee interaction patterns. Directed interactions (grooming, noncontact aggression, pant grunts/supplants) were consistent despite the inclusion of 18 months of data per community, indicating that most dyads interacted at relatively constant rates throughout the study period. Body contact and proximity showed lower consistency than directed interactions, most likely because a certain level of tolerance in foraging and resting extends to most group members, adding random noise that is not present in directed interactions. For body contact, no clear species-specific pattern emerged, but proximity (3m distance) was much less consistent in mangabeys than in chimpanzees, a result in line with recent findings regarding high levels of randomness in sooty mangabey spatial association patterns (Mielke et al., 2020). Just like rare interaction types, common but highly inconsistent interaction types could add noise to social relationship indices or when comparing network overlap.

While many animal species are studied in great detail, and vast amounts of long-term data are available, it is surprisingly difficult to convey the structure of social interactions across sites and species. Our consistency measure may help by providing a standardised way to convey the flexibility in interaction patterns over time and identify interaction types that likely differ in complexity between species. Further, many researchers use multilevel social network analysis and create relationship indices including different interaction types, unsure whether all of them will be equally reliable. This consistency measure, like similar efforts for hierarchies (Sánchez-Tójar et al., 2018), can be a useful tool to make these decisions while conveying important information about the study species. Importantly, these results further cement that researchers need to report sample sizes not only of their outcome variable, but also for interaction types that might have gone into creating relationship indices or network measures, because this gives readers the ability to judge the error associated with this predictor variable or network. To assess changes in relationships over time, there has been a trend to cut datasets into smaller subsets and then compare network overlap between these, assuming that the data in each are sufficient to depict the underlying distribution in the community. With our consistency measure, seasonality and change could be established if smaller subsets would show higher consistency than larger subsets, as random subsets retained consistent time intervals. This was not the case for any interaction type, even though some interaction types showed large variation, an indication that consistency is high during some times but not others.

Predictability is an important aspect of social complexity: an individual living in a system where all future social interactions are largely pre-defined by a few re-occurring factors needs little information to make decisions about its own behaviour (Flack, 2012; Sambrook & Whiten, 1997). Our consistency measure captures one aspect of predictability: if individuals distribute their social interactions the same across time, it is likely easy for group members to predict future social choices. This measure can easily be combined with other standardised approaches to social complexity and should mirror patterns (Fischer et al., 2017; Thierry et al., 2008). We did not find one consistent pattern of consistency difference between sooty mangabeys and chimpanzees; rather, variation within species was larger than between species, and each species showed higher consistency in some of the interaction types. One-dimensional measures of social complexity, such as group size, are thus probably insufficient to capture species differences in social complexity, as ‘complexity’ probably does not affects all aspects of life in a species uniformly: different species face different challenges, creating uncertainty in different areas of their social lives. Our consistency measure can detect which areas these are. Dyadic distributions of aggression and dominance interactions were highly predictable across groups. Spatial proximity was the least predictable aspect for all three groups, but as we have reported before, mangabey association beyond body contact contains large uncertainty (Mielke et al., 2020). Grooming interactions were less predictable in chimpanzees, indicating more varied grooming partner choice or changes over time. Many challenges are shared between primate species, especially regarding dyadic interaction patterns: It is therefore worth in a next step to consider the challenges arising from structuring interactions as sequences in time and the uncertainty arising when third parties influence decision making (Wittig et al., 2014). Our consistency measure offers a valuable piece in the puzzle of social complexity across animal species.

## Data Availability

Data and R scripts for the consistency analyses and simulations are available: https://github.com/AlexMielke1988/Mielke-et-al-Consistency

## Competing Interests

We have no competing interests

## Funding

AM, AP, LS, CC, RMW were supported by the Max Planck Society; AM was supported by the Wenner Gren Foundation (Grant Number 9095) and the British Academy Newton International Fellowship; AP was supported by the Leakey Foundation; LS was supported by the Minerva Foundation; JFG was supported by an NSF Graduate Research Fellowship (DGE-1142336), the Canadian Institutes of Health Research’s Strategic Training Initiative in Health Research’s Systems Biology Training Program, an NSERC Vanier Canada Graduate Scholarship (CGS), and a long-term Research Grant from the German Academic Exchange Service (DAAD-91525837-57048249). C. C. was supported by the European Research Council (ERC) under the European Union’s Horizon 2020 research and innovation programme (grant agreement no. 679787). RMW was supported by DFG Researcher Unit (FOR 2136) ‘Sociality and Health in Primates’ (WI 2637/3-1). Research at the Taï Chimpanzee Project has been funded by the Max Planck Society since 1997.

## Research Ethics

This study was purely observational with no manipulation of animals. Methods were approved by the Ethikrat der Max-Planck-Gesellschaft (4.08.2014).

## Permission to carry out fieldwork

Permissions to conduct the research were granted by the Ministries of Research and Environment of Ivory Coast (379/MESRS/GGRSIT/tm) and Office Ivorien des Parcs et Reserves.

## Acknowledgements

We thank the Ivorian Ministry of Environment and Forests and Ministry of Higher Education and Scientific Research and the Office Ivoirien des Parcs et Reserves of Côte d’Ivoire. We thank Simon Kannieu, Daniel Bouin, Gnimion Florent, Fabrice Blé, Florent Goulei, Apollinaire Gnahe Gjirian, Fredy Oulai Yehanon and the team of the TCP for field work support and data collection.

## Notes

https://github.com/AlexMielke1988/Mielke-et-al-Consistency

### Competing Interest Statement

The authors have declared no competing interest.

